# The Infraslow Fluctuation of Sigma Power During Sleep: Links to Markers of Arousal and Memory Reactivation Across Development

**DOI:** 10.1101/2024.11.06.620875

**Authors:** Maria E. Dimitriades, Alejandro Osorio-Forero, Sara Fattinger, Simone von Arx, Salome Kurth, Fiona Pugin, Valeria Jaramillo, Carina Volk, Elena Krugliakova, Melanie Furrer, Sven Leach, Peter Achermann, Miriam Gerstenberg, Reto Huber

## Abstract

Sleep is both a state of disconnection from the environment and a critical period for restoration. But how does sleep balance responsiveness with the protection of key functions? The infraslow fluctuation of sigma power (ISFS)—the clustering of sleep spindles over 10-100 seconds—is thought to regulate this trade-off in rodents. However, the organization of arousal and memory reactivation markers within the human ISFS and its conservation in younger ages remain unclear. This study characterizes the ISFS from childhood to young adulthood (N = 154; ages 8-26), examining its relationship with functional markers. Results indicate that the ISFS is present across all ages, with frequency, variability, and strength increasing from early to late adolescence. Notably, markers of arousal and memory reactivation are organized within the spindle-rich ISFS peak. The consistent presence and organization of the ISFS suggest it is intrinsic to sleep, with adolescence marking a dynamic window. These insights may guide interventions to promote healthier sleep across development.

## Introduction

Sleep comes with considerable evolutionary risks—reduced responsiveness and increased vulnerability to potential dangers. A growing hypothesis, based on rodent studies, suggests that sleep is not merely a vulnerable state but rather orchestrates a balance between environmental awareness and critical protected periods for learning and memory^1,2^. Recent research highlighted the importance of activity occurring on an infraslow time scale, over the course of 10 to 100 seconds, in achieving this balance^1,3^. In particular, sleep spindles—brief oscillatory bursts in the sigma range (10 to 16 Hz) that occur during non-rapid eye movement (NREM) sleep—do not occur uniformly throughout sleep, but instead fluctuate on an infraslow time scale^4^.

In rodents, the infraslow fluctuation of sigma power (ISFS) is driven by the locus coeruleus, a brainstem nucleus that regulates arousal through norepinephrine release^2^. The rodent ISFS segments NREM sleep into alternating fragile and protected periods^1,2,5^. Fragile periods demonstrate decreasing sleep spindle activity and increasing locus coeruleus activity, which is associated with a reduced arousal threshold in response to external stimuli. In contrast, protected periods show increasing sleep spindle activity, an elevated arousal threshold, and markers of memory reactivation. Researchers have interpreted this dynamic as an evolutionary trade-off, where fragile periods prioritize sensory vigilance and environmental awareness, while protected periods favor internal processing, including memory reactivation, by minimizing external disturbances^1^. Yet, the organization of markers of arousal and memory reactivation remain understudied in humans^6^.

While studies in adults have confirmed the presence of the ISFS in humans^1,6–8^, its trajectory and functional significance remain largely unexplored in child and adolescent populations. Childhood and adolescence represent critical periods of intense learning and brain maturation, marked by significant shifts in sleep structure, regulation, and electrophysiological characteristics, including sleep spindles, alongside a higher incidence of sleep problems, such as impaired sleep continuity^9–16^. Given the relationship between sleep, arousal, and memory reactivation during these developmental stages, examining the ISFS may provide insights into how these processes interact to support learning and cognitive development.

This research aims to analyze sleep electroencephalographic (EEG) data from children, early adolescents, late adolescents, and young adults to: 1) assess the presence and parameters of the ISFS from childhood to adulthood and 2) investigate how the ISFS relates to markers of arousal and memory reactivation throughout this developmental trajectory.

## Materials and Methods

### Participants

EEG sleep data were pooled from studies collected between 2008 and 2021^17–22^ conducted in the sleep laboratory of the University Children’s Hospital Zurich. Individuals were aged 8 to 26 years (N = 154, 16.3 ± 5.1 years, 38.3% female and 61.7% male) and four age groups were derived: children (8 to 11 years old; N = 39, 10.4 ± 1.1 years, 41.0% female and 59.0% male), early adolescents (12 to 15 years old; N = 45, 14.0 ± 1.2 years, 13.3% female and 86.7% male), late adolescents (16 to 20 years old; N = 32, 18.4 ± 1.7 years, 53.1% female and 46.9% male), and young adults (21 to 26 years old; N = 38, 23.5 ± 1.6 years, 55.3% female and 44.7% male). Individuals were excluded if they had a diagnosed sleep disorder, a history of mental health disorders, consumed psychotropic medications, and/or did not abide by a regular sleep schedule the week prior to the recording. Written informed consent and/or assent was obtained from all participants and/or legal guardians, respectively. Data collection was approved by the cantonal ethics committee, and all study procedures were performed according to the Declaration of Helsinki.

### Electroencephalographic Data

EEG sleep data were recorded using a 128-channel system (Electrical Geodesic Sensory Net; sampling rate: 500 or 1000 Hz). Sleep staging and artifact rejection were available from previous analyses^17–22^. Sleep staging was performed on 20 second epochs based on the American Association of Sleep Medicine criteria by two sleep experts^23^. Semi-automatic rejection of artifacts was performed based on power in two frequency bands (delta and beta) as described by Huber and colleagues^24^ and Leach and colleagues^25^. Only epochs that were artifact free in all channels were considered for further analyses. Excluded channels due to artifacts were interpolated from neighboring channels when 10% or less of the total channels were of poor quality. Next, EEG data were low-pass filtered (−3 dB cut-off: 39.25 Hz; Hamming windowed sinc Finite Impulse Response (FIR) filter from eeglab, filter order: 184), downsampled (200 Hz), and high-pass filtered (−3 dB cut-off = 0.27 Hz, Kaiser Window FIR filter, filter order: 2390). The data were re-referenced to the average of 109 electrodes; the electrodes on the edge of the net, which cover neck and face muscles, were excluded^25,26^.

### Analyses of the Infraslow Fluctuation of Sigma Power

Bouts of NREM Stage 2 (N2) sleep data that lasted for at least 280 seconds and were uninterrupted by artifacts or transitions to other vigilance states were isolated (demonstrated in **Figure 1A**). The minimum bout length of 280 seconds was chosen to ensure two full cycles of the lowest frequency of interest (0.0075 Hz). Our analyses focused on N2 sleep, where the ISFS is reported to be most prominent^1^. All subsequent analyses were performed on the aforementioned bouts.

**Figure 1:**
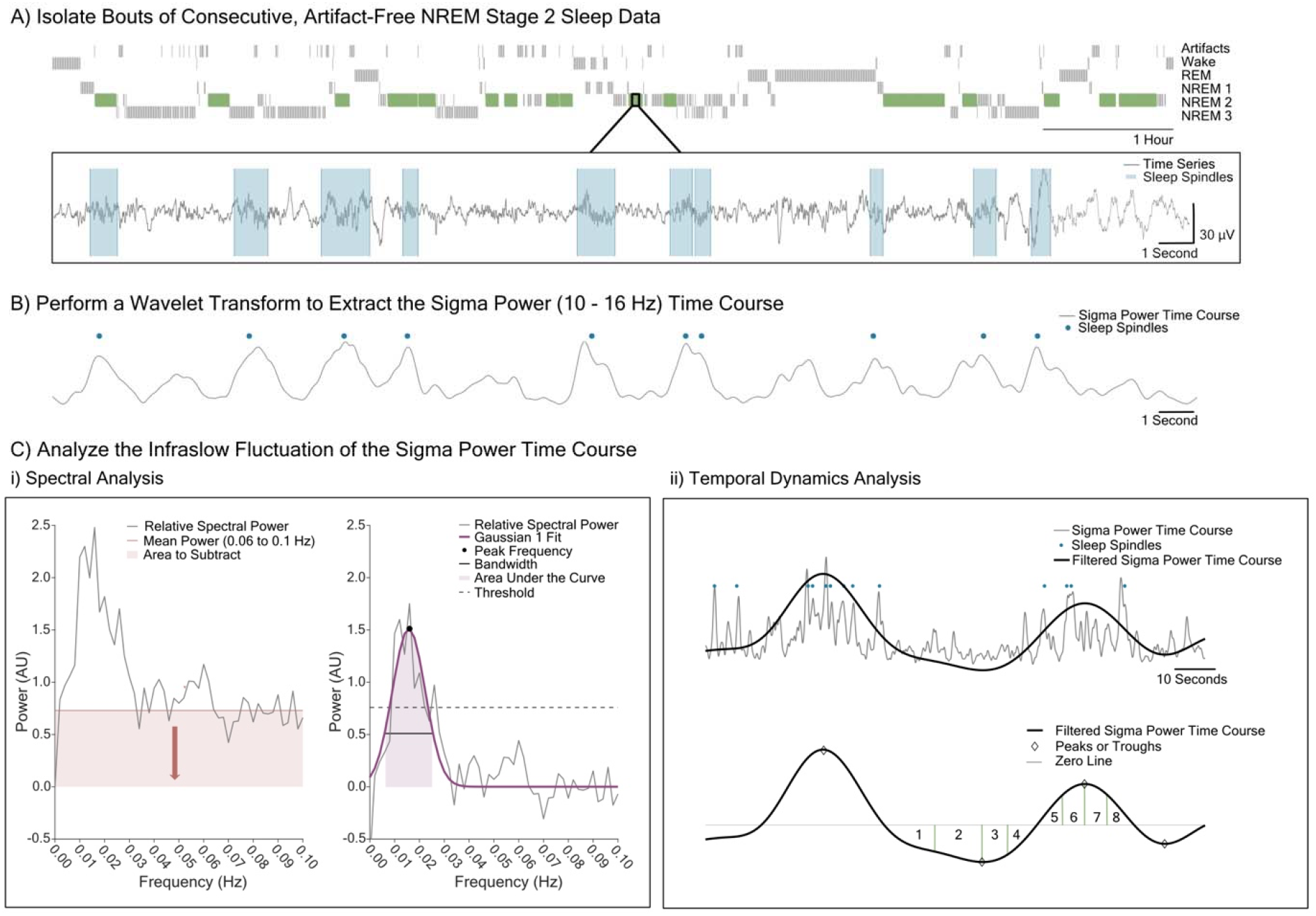
Overview of Analyses Assessing the Infraslow Fluctuation of Sigma Power. **A)** Top: Example hypnogram with bouts of consecutive, artifact-free non-rapid eye movement (NREM) stage 2 sleep lasting at least 280 seconds highlighted in green. Bout data will be used in all subsequent analyses. Bottom: Time series data is shown in grey with detected sleep spindles highlighted in blue. Sleep spindles are identified using an automatic detection method. **B)** A wavelet transform is performed on the time series data (corresponds to the data seen in A; sleep spindles marked as blue dots) to derive the sigma power time course (the mean of the absolute value of the sigma power time courses derived from 10 to 16 Hz) which is shown in grey. **C)** Outline of the two key analyses of the infraslow fluctuation of sigma power (ISFS) to be performed on the sigma power time course as presented in B: **i)** Left: The relative spectral power (the mean spectral power derived over all bouts for each channel and subject) is shown in grey. The relative spectral power between 0.06 to 0.1 Hz, indicated by a red horizontal line, is subtracted for baseline correction. Right: The resulting relative spectral power following baseline correction is shown as a grey line. A Gaussian function (Gaussian 1) is fit to the relative spectral power, shown in dark purple. If the peak power value surpasses the threshold, shown with a dotted line and calculated as 1.5 times the standard deviation of the gaussian, the presence of the ISFS is detected and parameters are further estimated: peak frequency (peak of Gaussian; marked with a black dot), bandwidth (±1 standard deviation from the peak frequency; black line), and area under the curve (total power within ±1 standard deviation from the peak frequency; shaded in light purple). **ii)** Top: The grey line represents the sigma power time course of part of one bout, with detected sleep spindles marked as blue dots (corresponding to B but over a longer time period to display the ISFS). The black line represents the filtered sigma power time course. Bottom: The filtered sigma power time course is standardized by subtracting the mean and dividing by the standard deviation before detecting peaks, troughs, and zero-crossings. The ISFS is further divided into eight phase bins; 1-4 denote the negative half wave and 5-8 denote the positive half wave. Any signal that could not be fully estimated, i.e., that did not occur on infraslow time scale (25 to 100 seconds) and/or was at the start or end of the bout and therefore did not contain both a peak and a trough, was labeled “Not ISFS”.

The sigma power time course was extracted from the N2 sleep bout data using a Mother Gabor-Morlet wavelet, following the method previously applied in mice^2^ (cycles: 4, frequency resolution: 0.2 Hz; demonstrated in **Figure 1B**). Considering that the frequency of sleep spindles increases across development and is generally higher in central compared to frontal regions^4,15^, we selected a broad frequency range, 10 to 16 Hz, to capture these differences in subsequent analyses. The absolute value of each sigma power time course derived from the bout, corresponding to the 31 input frequencies (ranging from 10 to 16 Hz in 0.2 Hz steps), was first calculated. The mean sigma power time course was then computed and used for subsequent analyses. To recenter the data for each bout around zero, the mean was subtracted from the sigma power time course.

#### Spectral Analysis of the Sigma Power Time Course

The Fast Fourier Transform (FFT) was applied to the sigma power time course of each bout (demonstrated in **Figure 1C**). To account for the varying lengths of the bouts (defined as lasting 280 seconds or more), the spectral power was adjusted by dividing the FFT output by the length of the corresponding bout. This adjustment ensures that the spectral power accurately corrects for the bout duration and allows for meaningful comparisons across different bouts. Some bouts exhibited a low-frequency component in the spectral power ranging from 0 to 0.004 Hz, which was not systematically linked to specific participants, channels, or sleep stages. This low-frequency component was occasionally removed with linear detrending and is believed to stem from the extraction of the sigma power time course. When the spectral power values in this low-frequency range exceeded the peak power detected outside this range (0.004 - 0.1 Hz), the power values from 0 to 0.004 Hz were excluded. This exclusion was applied before calculating the mean spectral power across all bouts, done separately for each channel and participant.

Relative power was calculated as the spectral power of each bout divided by the mean power within that same bout for each channel and participant; only power values from 0 to 0.1 Hz were considered. Absolute power was not reported in this project due to the higher EEG amplitude values typically observed in children and adolescents^17^, which significantly impact this measure. To detect a peak in the infraslow range and parameterize the curve, a Gaussian function (Gaussian 1) was fitted to the spectral power. Prior to fitting, the mean power value from 0.06 to 0.1 Hz was subtracted from the signal as a horizontal baseline. This correction was applied to remove any offset, as this frequency range visually tends to plateau before the spectrum rises to a peak near 0.25 Hz, as previously described by Achermann and Borbély^27^. Further, the frequency range of interest for detecting the peak of the ISFS was set between 0.01 and 0.04 Hz as this range typically contains the infraslow spectral peak^1,2^.

When the peak power value surpassed the threshold, defined as 1.5 times the standard deviation of the Gaussian curve, the signal in the given channel was considered to contain a detectable ISFS. The ISFS was characterized by three parameters: peak frequency (representing the most common period of the ISFS), bandwidth (representing the variability in period of the ISFS, calculated as ±1 standard deviation away from the peak frequency), and area under the curve (representing the strength of the ISFS, calculated as the total power within ±1 standard deviation away from the peak frequency). The bandwidth and area under the curve are defined by the same range of frequencies making these measures interrelated. To analyze age-related differences in the spectral parameters, the mean value across all electrodes was computed, excluding electrodes without a detectable ISFS. Additionally, to assess the topographical distribution of the ISFS spectral parameters independently of age-related changes, relative values were calculated by dividing the value of each electrode by the mean value of all electrodes for that participant. The spectral analysis of the ISFS was performed on the total and reduced number of bouts (see **Table 1**) to ensure comparability between the age groups.

**Table 1:** Parameters of Sleep Architecture. CH: Child group; EA: Early Adolescent group; LA: Late Adolescent group; YA: Young Adult group; SD: Standard Deviation; NREM: Non-Rapid Eye Movement (N1, N2, N3); REM: Rapid Eye Movement; min: minutes; s: seconds; %: percent. Bout: N2 sleep data that lasts for at least 280 seconds and is uninterrupted by artifacts or transitions to other vigilance states. Total Bout Sleep: The maximum number of bouts are taken from every individual. Reduced Bout Sleep: The last two bouts are excluded from the child, early adolescent, and late adolescent groups to make the bout parameters comparable. Mean Relative Location: the mean across all bout midpoints divided by the total length of the recording. Proportion of all N2: the total duration of bout data is divided by the total amount of N2 sleep. Corrected p-values and the direction of the difference between age groups is plotted with significantly different pvalues written in bold.

| Parameters | CH<br>Mean<br>(SD) | EA<br>Mean<br>(SD) | LA<br>Mean<br>(SD) | YA<br>Mean<br>(SD) | Corrected p-values and<br>Direction |
| --- | --- | --- | --- | --- | --- |
| **All Night Sleep** |  |  |  |  |  |
| N1<br>(min) | 27.39<br>(13.14) | 22.33<br>(13.42) | 27.03<br>(18.39) | 25.76<br>(12.23) | 0.24 |
| N2<br>(min) | 210.81<br>(45.43) | 213.38<br>(40.43) | 214.59<br>(32.35) | 224.32<br>(29.59) | 0.45 |
| N3<br>(min) | 139.62<br>(49.86) | 138.76<br>(42.87) | 85.93<br>(28.45) | 79.45<br>(32.04) | LA < CH (<0.0001)<br>YA < CH (<0.0001)<br>LA < EA (<0.0001)<br>YA < EA (<0.0001) |
| REM<br>(min) | 92.23<br>(23.31) | 83.74<br>(27.22) | 88.61<br>(30.28) | 98.14<br>(23.74) | 0.12 |
| Total Sleep<br>(min) | 470.05<br>(43.37) | 458.21<br>(52.66) | 416.12<br>(53.36) | 427.68<br>(30.96) | LA < CH (<0.0001)<br>YA < CH (<0.001)<br>LA < EA (<0.01)<br>YA < EA (0.01) |
| **Total Bout Sleep** |  |  |  |  |  |
| Bout Number | 11.36<br>(4.78) | 11.47<br>(4.47) | 11.97<br>(4.32) | 8.97<br>(4.43) | 0.05 |
| Mean Bout<br>Length (s) | 495.23<br>(95.24) | 470.68<br>(102.03) | 489.28<br>(65.72) | 458.39<br>(113.16) | 0.12 |
| Total Duration<br>of Bouts (min) | 97.59<br>(48.55) | 91.36<br>(43.21) | 99.23<br>(41.93) | 72.56<br>(43.33) | 0.07 |
| Mean Relative<br>Location (%) | 56.43<br>(7.30) | 56.02<br>(8.27) | 52.87<br>(6.58) | 49.36<br>(11.68) | YA < CH (0.02)<br>YA < EA (0.03) |
| Proportion of<br>all N2 | 44.10<br>(17.79) | 41.97<br>(16.88) | 45.69<br>(16.06) | 32.18<br>(18.87) | YA < CH (0.03)<br>YA < LA (0.01) |
| **Reduced Bout Sleep** |  |  |  |  |  |
| Bout Number | 9.51<br>(4.49) | 9.51<br>(4.38) | 9.97<br>(4.32) | 8.97<br>(4.43) | 0.96 |
| Mean Bout<br>Length (s) | 489.27<br>(89.10) | 467.63<br>(106.58) | 485.82<br>(70.97) | 458.39<br>(113.16) | 0.18 |
| Total Duration<br>of Bouts (min) | 80.79<br>(42.83) | 76.47<br>(41.62) | 82.22<br>(40.12) | 72.56<br>(43.33) | 0.78 |
| Mean Relative<br>Location (%) | 50.15<br>(7.85) | 49.44<br>(9.10) | 45.61<br>(7.76) | 49.36<br>(11.68) | 0.16 |
| Proportion of<br>all N2 | 36.24<br>(15.78) | 34.70<br>(16.60) | 37.63<br>(15.62) | 32.18<br>(18.87) | 0.55 |

#### Temporal Dynamics Analysis of the Sigma Power Time Course

The sigma power time course of each bout was also analyzed to study the ISFS over time in one central channel (C4; demonstrated in **Figure 1C**). The sigma power time course was low-pass filtered (−3 dB cut-off = 0.04 Hz, Butterworth Infinite Impulse Response (IIR) filter, filter order: 5 using filtfilt, Tukey window with a cosine fraction of 0.1) and then standardized by subtracting the mean and dividing by the standard deviation. All peaks, troughs, and zero-crossings (when the filtered sigma power time course crossed the zero-line corresponding to the mean of the signal) were identified. The presence of the ISFS was identified when two negative zero-crossings were separated by 25 to 100 seconds (i.e., 0.01 - 0.04 Hz, corresponding to our frequency range of interest mentioned above).

To assess where electrophysiological characteristics (defined below) occurred within the ISFS, the filtered sigma power time course defining the ISFS was divided into eight phase bins. The start of the ISFS (first negative zero-crossing) until the midpoint (positive zero-crossing) is the negative half wave of the ISFS. It is further divided by the trough: the mean sample between the start and the trough separates bins 1 and 2 and the mean sample between the trough and the midpoint separates bins 3 and 4. Moreover, the midpoint of the ISFS until the end of the ISFS (second negative zero-crossing) is the positive half wave of the ISFS. It is further divided by the peak: the mean sample between the midpoint and the peak separates bins 5 and 6 and the mean sample between the peak and the end separates bins 7 and 8. Any signal that could not be fully estimated, i.e., that did not occur on infraslow time scale (25 to 100 seconds) and/or was at the start or end of the bout and therefore did not contain both a peak and a trough, was labeled “Not ISFS”.

### Detection of Electrophysiological Characteristics

No systematic protocol was established across studies to evoke arousals, resulting in a limited number of spontaneous full arousals. Consequently, we concentrated on detecting spontaneous microarousals by adapting an established algorithm^28,29^ to capture a broader range of responses, including shorter responses starting from 1.5 up to 10 seconds^30–32^. In brief, this algorithm uses EEG and/or electromyographic data to detect arousals by identifying increases in baseline alpha and/or beta activity above a relative threshold, signaling the presence of an arousal event. To ensure that microarousals detected using alpha-band criteria did not overlap with sleep spindles—especially in younger individuals with typically lower sleep spindle frequencies^15^—each microarousal was visually inspected to ensure it was not a sleep spindle. Further, we repeated the analysis examining the distribution of arousals within the ISFS 1) using only beta-band arousals to avoid any overlap with spindles and 2) with the original algorithm assessing full arousals to ensure that while fewer full arousals were detected, the overall pattern within the ISFS remained consistent. The onset of the given microarousal or arousal was taken as the sample of the event.

Sleep spindles were identified using an established amplitude-threshold algorithm^33^. In short, N2 sleep bout data were isolated and band-pass filtered (−3 dB cut-offs: 9.96 and 17.5 Hz; Chebyshev Type II IIR filter; filter order: 6 using filtfilt). Given the variability in signal amplitude across electrodes, relative upper and lower thresholds were established for each channel and participant. Sleep spindles were detected in a given channel when the amplitude exceeded an upper threshold of five times the average signal amplitude and until the instances where the signal surrounding the maximum peak fell below two times the average signal amplitude for the given channel. The minimal duration of a sleep spindle was set to be 0.3 s^34^.

Slow waves were detected using an established algorithm^35^. Bouts of N2 sleep data were band-pass filtered (−3 dB cut-offs: 0.34 and 7.1 Hz; Chebyshev Type II IIR filter; filter order: 2 using filtfilt) and all peaks, troughs, and zero-crossings were identified. Slow waves were identified as the time between two negative zero-crossings, separated by intervals ranging from 0.5 to 2 seconds. To exclude outliers, the bottom 10% and top 1% of peak-to-trough amplitudes were removed. Further, in order to target high-amplitude slow waves, likely representing K-complexes, only those within the top 80th percentile of peak-to-trough amplitudes were considered for analysis as previously reported^35^. Slow wave-sleep spindle coupling was further defined when the maximum amplitude of a detected sleep spindle fell within the up-phase of a detected slow wave.

To assess the distribution of microarousal onsets within each phase bin of the ISFS, the number of detected microarousals in each phase bin of the ISFS was calculated as a percentage of the total number of microarousals detected for each participant; only individuals with at least three microarousals were included. The same calculation was done to assess the distribution of sleep spindles, slow waves, and slow wave-sleep spindle coupling within the phase bins of the ISFS. The relative coupling of slow waves and sleep spindles was also quantified by dividing the number of sleep spindles coupled to a slow wave within a given phase bin of the ISFS by the total number of sleep spindles in that phase bin of the ISFS.

### Statistical Analysis

All statistical analyses were conducted in MATLAB R2022a. Normality was assessed using quantile-quantile plots and Shapiro-Wilk tests. Kruskal-Wallis tests, followed by Tukey-Kramer post-hoc tests, were used to compare sleep architecture parameters and the number of microarousals occurring during N2 bout data for the four age groups and the distribution of microarousals, sleep spindles, slow waves, and slow wave-sleep spindle coupling within the phase bins of the ISFS. The same was done to assess the temporal occurrence of sleep spindles around the onset of microarousals, where only individuals who had at least three sleep spindles after the onset of microarousals were included. Mann-Whitney U tests with false discovery rate (Benjamini-Hochberg) correction were conducted to compare the percentage of sleep spindles coupling within slow waves across conditions. One-way ANOVAs were used to compare the ISFS spectral parameters across channels, with Tukey-Kramer post-hoc tests for multiple comparisons. Electrode-wise topographical differences were assessed using one-way ANOVAs, followed by cluster-based permutation testing. A p-value less than 0.05 was considered significant.

## Results

### Sleep Architecture Parameters

Sleep architecture parameters have been previously published^17–22^. All-night sleep parameters showed that the total amount of NREM sleep stage N1 and N2 and Rapid Eye Movement (REM) sleep were comparable between groups (**Table 1**). As expected, a significant increase in N3 sleep and total sleep time was detected in the child and early adolescent groups when compared to the late adolescent and young adult groups.

To analyze data related to the ISFS, continuous N2 sleep bout data were examined. The number of bouts, mean bout length, and the total duration of N2 sleep bout data were similar across age groups.

### The Presence and Spectral Parameters of the Infraslow Fluctuation of Sigma Power

Spectral analysis of the sigma power time course revealed that the ISFS was detectable in children, early adolescents, late adolescents, and young adults (**Figure 2A**). All individuals had a detectable ISFS in at least 20% of electrodes (mean +/-standard deviation in % of electrodes with a detectable ISFS: children: 74.74 +/-21.08, early adolescents: 75.66 +/-21.83, late adolescents: 82.05 +/-18.14, young adults: 80.87 +/-24.85). The mean value of each ISFS spectral parameter—peak frequency, bandwidth, and area under the curve—was calculated across channels. All ISFS spectral parameters showed significant differences between age groups (**Figure 2B**): peak frequency was significantly higher in the young adult group compared to the child and early adolescent groups (percent change: children vs. young adults = 22.37, early adolescents vs. young adults = 17.24; F(3,150) = 4.60, p = 0.004), bandwidth was significantly higher in the late adolescent group compared to the child group (percent change: children vs. late adolescents = 52.59; F(3,150) = 6.32, p < 0.001), and area under the curve was significantly higher in the late adolescent and young adult groups compared to the child group (percent change: children vs. late adolescents = 45.22, children vs. young adults = 38.47; F(3,150) = 5.38, p = 0.002).

**Figure 2:**
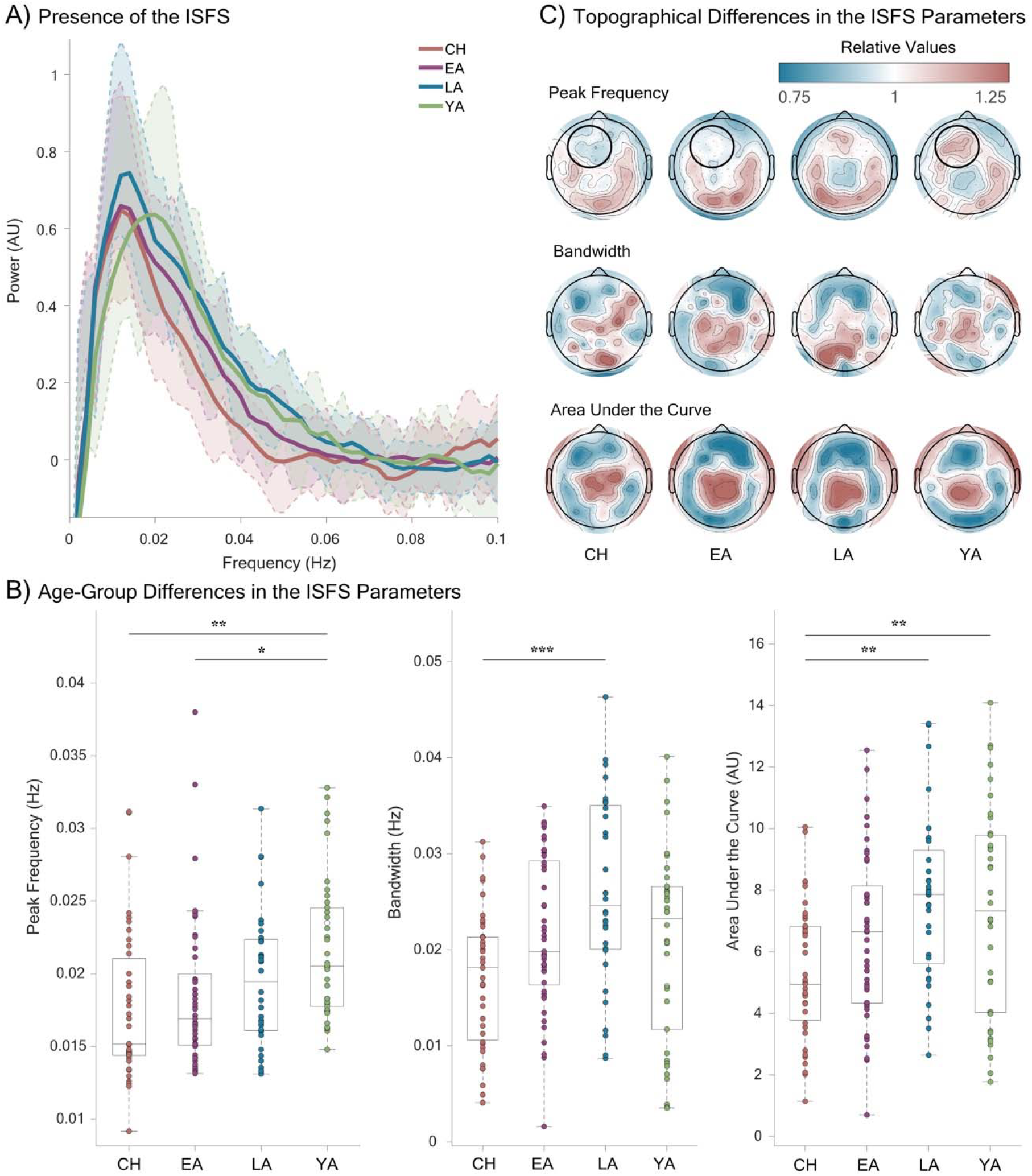
The Presence and Spectral Parameters of the Infraslow Fluctuation of Sigma Power Across Development. **A)** Bold lines represent the smoothed mean spectral power of the infraslow fluctuation of sigma power (ISFS) averaged across all electrodes for each age group (CH: children; EA: early adolescents; LA: late adolescents; YA: young adults). The standard deviation for each age group is shown in paler colors with dotted lines. **B)** The mean spectral parameters of the ISFS (peak frequency, bandwidth, area under the curve) averaged across all electrodes are shown for individuals (denoted by dots) for all age groups. Significant differences between age groups are indicated by an asterisk(s). **C)** Rows represent the relative values of each spectral parameter of the ISFS, while columns correspond to different age groups. A circle in the top row highlights a cluster of frontal electrodes that show significant differences across the age groups.

Topographical analysis of the ISFS spectral parameters found that most electrodes showed no significant differences across age groups (**Figure 2C**). However, regarding peak frequency, a frontal cluster of seven electrodes revealed significantly higher values in the young adult group compared to the child and early adolescent groups (cluster mean +/-standard deviation: children = 0.94 +/-0.02, early adolescents 0.98 +/-0.01, young adults 1.11 +/-0.04; young adult vs. child p = 0.009; child vs. young adult p = 0.027), indicating higher peak frequencies frontally with age. Peak frequency and area under the curve showed local minima and maxima in central regions, respectively, while bandwidth displayed no clear topographical pattern.

All the above analyses could be confounded by significant age-related differences in the relative location of each bout (within the total recording) and the proportion of N2 bout data (relative to total N2 sleep; **Table 1**). Thus, in order to compare a similar distribution and proportion of bout sleep data, the two final bouts were removed from the child, early adolescent, and late adolescent groups, which resulted in no statistical difference between groups for the bout parameters (**Table 1**). The repeated spectral analyses revealed similar age- and topographical-related changes of the ISFS spectral parameters (data not shown).

### Organization of Electrophysiological Characteristics within the Infraslow Fluctuation of Sigma Power

All subsequent analyses focus on microarousals, sleep spindles, slow waves, and slow wave-sleep spindle coupling in a central electrode, chosen for its robust ISFS across all age groups as indicated by a high area under the curve (reported in **Figure 2C**). To assess the organization of electrophysiological characteristics within the ISFS, the ISFS was divided into eight phase bins. The amount of data in the positive and negative half waves of the ISFS was comparable (49-51%), as well as the amount of data in each of the eight phase bins of the ISFS and the data where the ISFS was not able to be estimated (Phase bins 1-8: 10-11%; Not ISFS: 12-15%) across age groups. This indicates that the distribution of electrophysiological characteristics across each half wave, phase bin, or Not ISFS was not biased by the duration of the time frame analyzed.

When investigating the distribution of the onset of microarousals within the ISFS, only participants with at least three detected microarousals were included (children = 22; early adolescents = 29; late adolescents = 22; young adults = 20). The microarousal index in the N2 bout data was similar across age groups (4-7 per hour; H(3) = [4.20]; p = 0.24); the mean duration of a microarousal was 3.94 seconds. The distribution of microarousals significantly differed between phase bins of the ISFS for all age groups (**Figure 3A**; children: H(8) = [36.09], p < 0.0001; early adolescents: H(8) = [17.21], p = 0.028; late adolescents: H(8) = [40.07], p < 0.0001; young adults: H(8) = [37.25], p < 0.0001). The highest percentage of microarousals occurred around the peak of the ISFS (corresponding to phase bin 6 and/or 7) compared to phase bin(s) occurring in the negative half wave of the ISFS for all age groups. This analysis was repeated using only beta band criteria for arousals and the original algorithm analyzing full arousals. Both additional analyses revealed the same distribution of arousals, with the majority occurring at the peak of the ISFS (data not shown).

**Figure 3:**
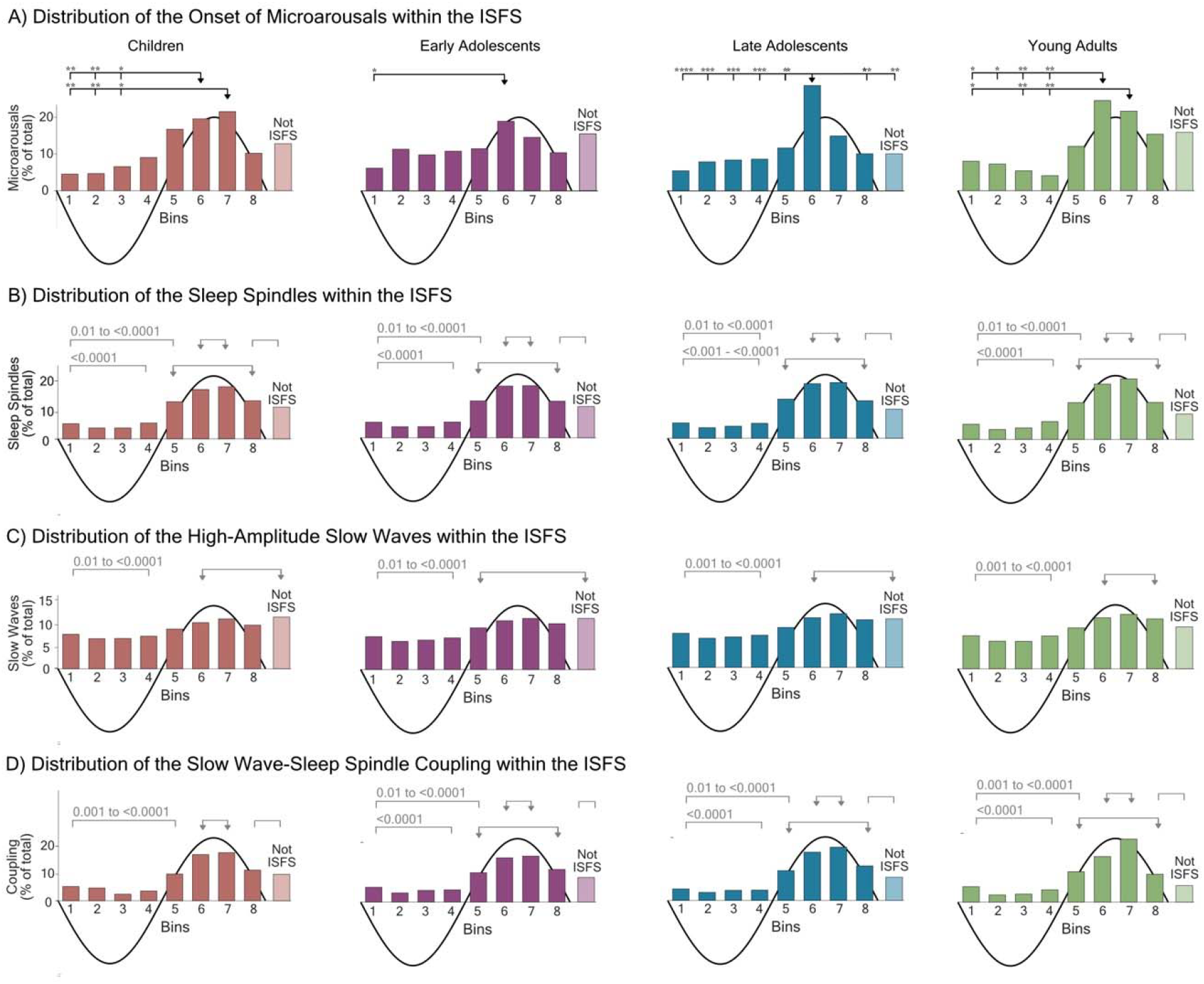
Distribution of Electrophysiological Characteristics within the Infraslow Fluctuation of Sigma Power. **A)** The distribution of the onset of microarousals across bins of the infraslow fluctuation of sigma power (ISFS) and in periods where the ISFS could not be estimated (denoted as “Not ISFS”) is displayed. Participants are excluded from this analysis if they did not have at least three microarousals. The black arrow indicates the phase bin(s) that is significantly higher when compared to other phase bins (denoted by black ticks). **B)** The distribution of sleep spindles across phase bins of the ISFS and Not ISFS is displayed. **C)** The distribution of high-amplitude slow waves across phase bins of the ISFS and Not ISFS is displayed. **D)** The distribution of slow wave-sleep spindle coupling across phase bins of the ISFS and Not ISFS is displayed. All electrophysiological characteristics are calculated by dividing the number of electrophysiological characteristics in each phase bin of the ISFS by the total number of electrophysiological characteristics for each participant, with the mean across participants displayed. Each column represents an age group, with significant differences between phase bins marked by asterisk(s). For groups and parameters with many significant differences between groups, grey arrows are used to show groups of phase bins that are significantly higher when compared to other phase bins (denoted in grey brackets); the range of p-values written above them in grey. The bold black wave illustrates the relative position of each phase bin within the ISFS.

The distribution of sleep spindles, slow waves, and slow wave-sleep spindle coupling also differed between phase bins of the ISFS for most age groups. Per definition, sleep spindles occurred most in the positive half wave of the ISFS (corresponding to phase bins 5 - 8; peak: 6 - 7) when compared to phase bins of the negative half wave of the ISFS (corresponding to phase bins 1 - 4; trough: 2 - 3) for all age groups (**Figure 3B;** children: H(8) = [287.78]; early adolescents: H(8) = [330.73]; late adolescents: H(8) = [242.13]; young adults: H(8) = [266.37]; all p-values were <0.0001). Within the ISFS, more high-amplitude slow waves occurred in and around the peak compared to the negative half wave for all age groups (**Figure 3C**; children: H(8) = [153.99]; early adolescents: H(8) = [210.28]; late adolescents: H(8) = [149.99]; young adults: H(8 = [155.58]; all p-values were <0.0001). Subsequently, the coupling of slow waves with sleep spindles also occurred more in the peak of the ISFS when compared to phase bins in the negative half wave of the ISFS for all age groups (**Figure 3D**; children: H(8) = [177.96]; early adolescents: H(8) = [211.83]; late adolescents: H(8) = [173.42]; young adults: H(8) = [147.57]; all p-values were <0.0001). Additionally, the relative coupling, i.e., the percent of sleep spindles that coupled with slow waves in a given phase bin of the ISFS after dividing by the total number of sleep spindles in the given phase bin, revealed that more coupling occurred in the peak of the ISFS when compared to the trough of the ISFS for the child and young adult groups (children: H(8) = [27.09], p = <0.001; young adults: H(8) = [15.7], p = 0.04); a trend level difference was found for the late adolescent group (H(8) = [13.70], p = 0.09). In summary, sleep spindles, slow waves, and slow wave-sleep spindle coupling were more likely to occur in the positive half wave of the ISFS when compared to the negative half wave of the ISFS.

### Sleep Spindle Occurrence and Slow Wave-Sleep Spindle Coupling following the Onset of Microarousals

To determine if sleep spindles temporally occur around the onset of microarousals, sleep spindle occurrence was compared in the five seconds before and after the onset of any detected microarousal (**Figure 4A**). Significant differences were found across all age groups (children: H(9,220) = [64.38], p < 0.0001; early adolescents: F(9,290) = [98.23], p < 0.0001; late adolescents: F(9,220) = [46.53], p < 0.0001; young adults: F(9,200) = [77.03], p < 0.0001). In all age groups, sleep spindles occurred more frequently one to two seconds after the onset of a microarousal compared to the five seconds before the onset of the microarousals and occasionally when compared to the three to five seconds after the onset of the microarousals.

**Figure 4:**
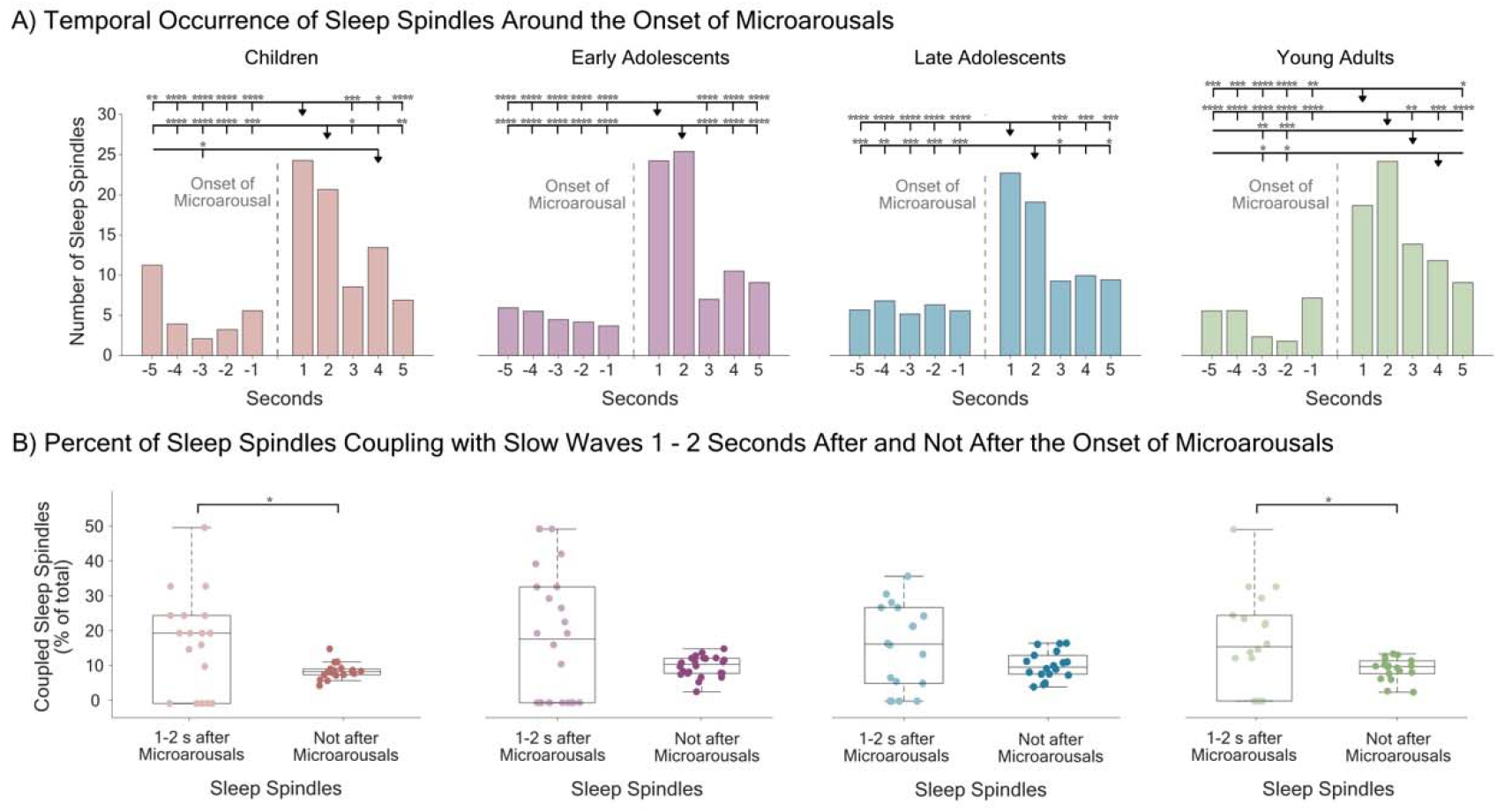
Relationship between Microarousals, Sleep Spindle Temporal Occurrence, and Slow Wave-Sleep Spindle Coupling Independent of the Infraslow Fluctuation of Sigma Power. **A)** The average temporal occurrence of sleep spindles across participants in the five seconds prior and five seconds post the onset of a microarousal is displayed. **B)** The number of spindles that couple into the up-phase of high-amplitude slow waves divided by the total number of spindles in the established time window, i.e., either in the one to two seconds after the onset of the microarousal (left) or any sleep spindle not occurring within this two second window (right) is displayed. Each dot denotes a participant; participants are excluded from this analysis if they did not have at least three microarousals and three sleep spindles following the onset of microarousals. Significant differences are denoted with asterisk(s).

The sleep spindles that occurred within the two seconds after the onset of microarousals were compared to all other sleep spindles in terms of coupling with slow waves (**Figure 4B**). Only individuals who had at least three microarousals and three sleep spindles following microarousals were included (children = 22; early adolescents = 29; late adolescents = 22; young adults = 20) in these analyses. The percentage of sleep spindles coupling with slow waves was significantly higher for spindles occurring one to two seconds after the onset of microarousals when compared to all other spindles in children (z = 2.14; p = 0.03) and young adults (z = 2.08; p = 0.03).

## Discussion

Our findings demonstrate that sigma power during N2 sleep fluctuates on an infraslow time scale from childhood to young adulthood, suggesting its fundamental biological and functional relevance. Yet, we observe that the frequency, variability, and strength of the ISFS increase from children and early adolescents to late adolescents and young adults. Notably, all age groups show a consistent organization in markers of arousal and memory reactivation, which occur the most in the spindle-rich peak of the ISFS and further support that the ISFS is intrinsic to sleep.

To optimize and systematically capture age-specific aspects, we refined previous methods for analyzing the ISFS in humans^1,8^. These adaptations included extending bout lengths to cover at least two cycles of our lowest frequency of interest (0.0075 Hz), removing filtering and downsampling steps that may reduce the temporal resolution, utilizing FFT instead of wavelets for the spectral analysis, and revising criteria for detecting and quantifying the ISFS. Despite these methodological changes, the mean peak frequency detected in the young adult group (0.21 Hz) remained consistent with previously reported values (0.2 Hz) for this age range^1^, supporting the validity of our approach.

Further analysis of the ISFS spectral parameters revealed significant developmental changes. Younger age groups exhibited lower peak frequency, indicating a longer average ISFS period, as well as narrower bandwidth, suggesting less variation in the ISFS period, and reduced area under the curve, reflecting less strength of the ISFS compared to older individuals. A recent nap study^15^ established age-specific normative values for sleep spindle characteristics, showing that spindle density increased from age three until about age 15, after which it plateaus into young adulthood. This aligns with our observation that ISFS parameters differ most significantly between younger (8 to 15 years old) and older (16 to 26 years old) groups. Given that the thalamus and cortex, the key regions involved in sleep spindle generation, undergo significant maturation during this period^9,11,14^, it is likely that these structural changes contribute to the developmental differences we observed in the ISFS parameters.

However, the presence of the ISFS was observed across all age groups, and the investigation of topographical patterns revealed that the ISFS spectral parameters exhibit largely similar spatial distributions, suggesting the absence of a clear developmental gradient in the spatial organization and emphasizing the fundamental nature of this rhythm. Therefore, despite ongoing cortical and subcortical maturation of underlying systems^11,36,37^, the anatomical structures facilitating these infraslow dynamics appear to be established early in life. Only peak frequency was found to increase with age in a frontal cluster, which may reflect the prolonged maturation of the frontal cortex during childhood and adolescence^38–40^.

While the thalamus and cortex are crucial for the generation and propagation of sleep spindles, the locus coeruleus is necessary and sufficient for the ISFS in rodents^2,4,41^. Analysis of the temporal dynamics of the ISFS has shown that when the ISFS is decreasing, norepinephrine is increasing, marking the fragile phase of the ISFS. Correspondingly, both around the peak of the ISFS and in the subsequent decreasing phase of the ISFS, external tones were more likely to induce wakefulness or microarousals (lasting 4 to 17 seconds and involving movement) in mice^1,42,43^. In contrast, decreasing locus coeruleus activity marks the protected phase of the ISFS, corresponding to around the trough and subsequent increasing phase of ISFS, which was associated with greater resilience to external stimuli and hippocampal ripple activity—an indicator of memory reactivation^1,5,44^.

Expanding this to humans, we found that the onset of spontaneous microarousals (lasting 1.5 to 10 seconds and not necessarily involving movement) were not randomly distributed within the ISFS. Instead, they clustered around the peak of the ISFS in all age groups. Thus, the organization of arousal within the human ISFS shows some overlap with rodent findings, though it does not extend as far into the decreasing phase of the ISFS as observed in rodents. In addition to microarousals, we observed that the percentage of high-amplitude slow waves and sleep spindles coupling with these slow waves—considered a marker of memory reactivation within the active-systems consolidation model^45^—was also higher during the peak of the ISFS. Overall, these findings support a temporal shift between the species, potentially driven by underlying anatomical or physiological factors, as previously speculated^1,6,46^. However, the variability in markers of arousal and memory reactivation and the absence of perturbation studies examining the ISFS in humans hinders cross-species comparisons.

Interestingly, microarousals were followed by an increase in sleep spindles, which may reflect the previously proposed protective role of spindles for sleep: individuals who generate more spindles demonstrate higher noise tolerance and promoting spindles pharmacologically or genetically has been shown to elevate the arousal threshold^47–49^. Moreover, we observed that in children and young adults, post-microarousal spindles were more likely to couple with slow waves. Taken together, a possible binding factor between spindles, microarousals, and memory reactivation could be K-complexes, which are often associated with spindles and play a role in regulating both arousability and memory reactivation and have been previously reported to occur most in the peak of the ISFS^6,22,50^.

While functional markers show a clear organization within the ISFS that is maintained across development, spectral parameters of the ISFS show pronounced differences between childhood/early adolescence and late adolescence/young adulthood. Whether these differences contribute to increased difficulties with sleep initiation and maintenance, commonly reported in children and adolescent populations^10,13^, remains to be determined.

Our study used retrospective datasets not specifically designed to perturb the ISFS or systematically assess overnight memory performance. Therefore, we only captured spontaneous arousal events, likely triggered by internal or random environmental factors, and memory reactivation inferred through electrophysiological markers. Further research exploring perturbations of the ISFS through external stimuli and its implications in various clinical populations and memory types will be crucial for understanding how these mechanisms shape sleep and memory throughout the lifespan. Additionally, the ISFS was measured indirectly with surface EEG, making it difficult to infer both the precise temporal dynamics and underlying anatomical sources of the ISFS.

In summary, our analyses demonstrate that sigma power fluctuates on an infraslow time scale across development, revealing age-specific differences in the ISFS spectral parameters. The increase in the frequency, variability, and strength of the ISFS across development may drive essential changes in the regulation of sleep and its relationship with memory reactivation.

Markedly, we find that readouts of arousal and memory reactivation are organized within the spindle-rich peak of the ISFS irrespective of age group. By elucidating how key sleep processes are organized within the ISFS across development, this study lays the groundwork for future investigations into the role of ISFS in shaping cognitive development and arousal regulation, with implications for understanding both typical and disrupted sleep patterns from childhood to adulthood.

## Data and Materials Availability

All data supporting the findings of this study are securely stored at servers of the University Children’s Hospital. Access and availability will be provided upon a material transfer agreement and after approval by the local ethics committee of the Canton of Zurich.

## Author Contributions

Data Collection: SK, FP, VJ, CV, EK, MF, SL

Methodology: MED, AOF, SF, SV, PA, MG, RH

Formal Analysis: MED

Supervision: MG, RH

Writing - Original Draft: MED

Writing - Editing: MED, AOF, SF, SL, PA, MG, RH

Writing - Reviewing: MED, AOF, SF, SA, SK, FP, VJ, CV, EK, MF, SL, PA, MG, RH

## Acknowledgements

We would like to thank the participants for their time and data contributions, as well as their families for their support. We also appreciate the invaluable advice and support from everyone in Reto Huber’s lab and Miriam Gerstenberg’s research group throughout this project.

